# The iron-sulfur cluster is critical for DNA binding by human DNA polymerase ε

**DOI:** 10.1101/2022.05.05.490830

**Authors:** Alisa E. Lisova, Andrey G. Baranovskiy, Lucia M. Morstadt, Nigar D. Babayeva, Elena I. Stepchenkova, Tahir H. Tahirov

## Abstract

DNA polymerase ε (Polε) is a key enzyme for DNA replication in eukaryotes. It is attached to a helicase and performs DNA synthesis on the leading strand. Recently it was shown that the catalytic domain of yeast Polε (Polε_CD_) contains a [4Fe-4S] cluster located at the base of the processivity domain (P-domain) and coordinated by four conserved cysteines. In this work, we have shown that human Polε_CD_ (hPolε_CD_) expressed in bacterial cells also contains an iron-sulfur cluster. In comparison, recombinant hPolε_CD_ produced in insect cells contains an eight-fold-lower level of iron. Interestingly, the iron content correlates with the level of DNA-binding molecules, which suggests an important role of the iron-sulfur cluster in hPolε interaction with DNA. Indeed, mutation of two conserved cysteines that coordinate the cluster abolished template:primer binding and, therefore, DNA polymerase and proofreading exonuclease activities. We propose that the cluster regulates the conformation of the P-domain, which, like a gatekeeper, controls access to a DNA-binding cleft for a template:primer. In addition, we performed kinetic and binding studies of hPolε_CD_. The binding studies demonstrated low affinity of hPolε_CD_ to DNA and a strong effect of salt concentration on stability of the hPolε_CD_/DNA complex. Pre-steady-state kinetic studies have shown a maximal polymerization rate constant of 51.5 s^-1^ and a relatively low affinity to incoming dNTP with an apparent *K*_*D*_ of 105 μM. This work provides notable insight into the role of a [4Fe-4S] cluster in Polε function.

## INTRODUCTION

DNA polymerase ε (Polε) is one of the main eukaryotic replicases responsible for leading-strand synthesis during genome duplication (1-3). It belongs to the B-family of DNA polymerases, which also contains Polδ, Polα, and Polζ. Human Polε (hPolε) consists of four subunits: the catalytic subunit (p261) and the accessory subunits p59, p17, and p12, listed in order from bigger to smaller. p261 is composed of two duplicated exonuclease/polymerase domains, the first of which is active, while the second is inactive and plays an important structural role (4-6). In the recently reported structure of yeast Polε holoenzyme, the small subunits p17 and p12 are tethering the two lobes of p261 (7). Instability of hPolε holoenzyme results in replication stress, tumorigenesis, and developmental abnormalities (8,9).

The Polε catalytic subunit contains three cysteine motifs: CysA, CysB, and CysX (10,11). The first two are located at the extreme C-terminus, which interacts with a B-subunit (p59 in humans). The recently discovered CysX motif is located in the N-terminal domain responsible for DNA polymerase and exonuclease activities. The first report about the presence of a [4Fe-4S] cluster in yeast Polε is dated to 2011, when the cluster location was attributed to a CysB motif (12). This cluster position was not confirmed in subsequent work by our group (13), which is consistent with the crystal structure of the human Polε subcomplex p59-p261(2142-2286) showing zinc in CysA and CysB (14). In 2014, Jain et al. discovered the CysX motif and demonstrated its importance for [4Fe-4S] coordination and DNA polymerase activity of yeast Polε (10). Later, the presence of a iron-sulfur cluster in the CysX motif of yeast Polε was confirmed using a structural approach (11). Recently it was shown that the iron-sulfur cluster of yeast Polε is redox active and its oxidation affects DNA polymerase activity (15). The role of the cluster in Polε interaction with DNA has not been studied so far.

In this work, we established that the [4Fe-4S] cluster is critical for template:primer binding by human Polε. Two point mutations that disrupt cluster coordination abrogate hPolε interaction with DNA, resulting in the loss of DNA polymerase and exonuclease activities. We also analyzed the outcome of the protein expression system on the level of iron and DNA-binding properties of hPolε samples. In addition, we conducted pre-steady-state kinetic and binding studies of hPolε at near-physiological salt concentration.

## RESULTS

### The iron-sulfur cluster is important for hPolε interaction with DNA

The catalytic domain of human DNA polymerase ε (hPolε_CD_; residues 28-1194) was expressed in *Escherichia coli* and purified to near homogeneity (Figure 1). The concentrated sample displayed a brownish color, which is characteristic of proteins containing an [4Fe-4S] cluster (10,11). Interestingly, the same protein overexpressed in insect cells and purified in the same way (purity is shown in Figure 1) was almost colorless. Analysis of the iron content revealed that the hPolε_CD_ sample expressed in insect cells has an 8.6-fold-lower iron level than the one expressed in *E. coli* (Table 1). This result indicates that insect cells cannot efficiently incorporate the iron-sulfur cluster into a heterologous protein during its overexpression.

**Table 1.** DNA-binding activity of hPolε_CD_ samples correlates with their iron content.

| Expression system | Iron content <sup>a</sup><br>% | Active molecules <sup>b</sup><br>% |
| --- | --- | --- |
| insect | 7.3 ± 0.93 | 8.3 ± 0.7 |
| *E. coli* | 63 ± 7.1 | 51.2 ± 4.1 |

<sup>a</sup> The iron content of four atoms per protein molecule was taken for 100%.
<sup>b</sup> The molecules showing DNA binding activity.
Data are presented as mean ± SD.

**Figure 1.**
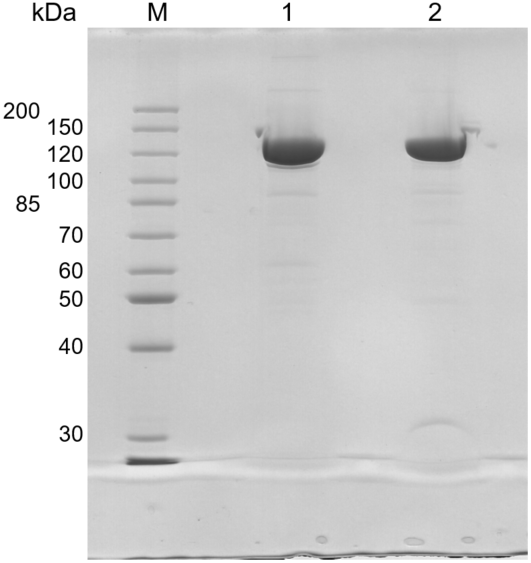
Analysis of purity of hPolε_CD_ samples. Samples were separated by 8% SDS-PAGE and stained by Coomassie Brilliant Blue R-250. M – markers; lane 1 - hPolε_CD_(insect); lane 2 - hPolε_CD_ (*E*.*coli*).

For both hPolε_CD_ samples, we analyzed the level of molecules possessing DNA-binding activity by using an electrophoretic mobility shift assay (EMSA; Figure 2). All reactions contained varying concentrations of hPolε_CD_ and 0.5 μM DNA duplex composed of a 15-mer primer annealed to a 25-mer template (Table 2). After 5 min of incubation at room temperature, the products were resolved by 5% native gel. This approach allows to separate DNA molecules from protein/DNA complexes and calculate the percentage of complexed DNA at each protein concentration. We found that hPolε_CD_ samples obtained after expression in *E. coli* and insect cells have 51% and 8.5% of active molecules, respectively (Table 1). The correlation between iron content and the level of DNA-binding molecules suggests that the iron-sulfur cluster is important for interaction of hPolε with DNA. Of note, all data described below were obtained using the active enzyme concentration if not otherwise indicated.

**Table 2.**
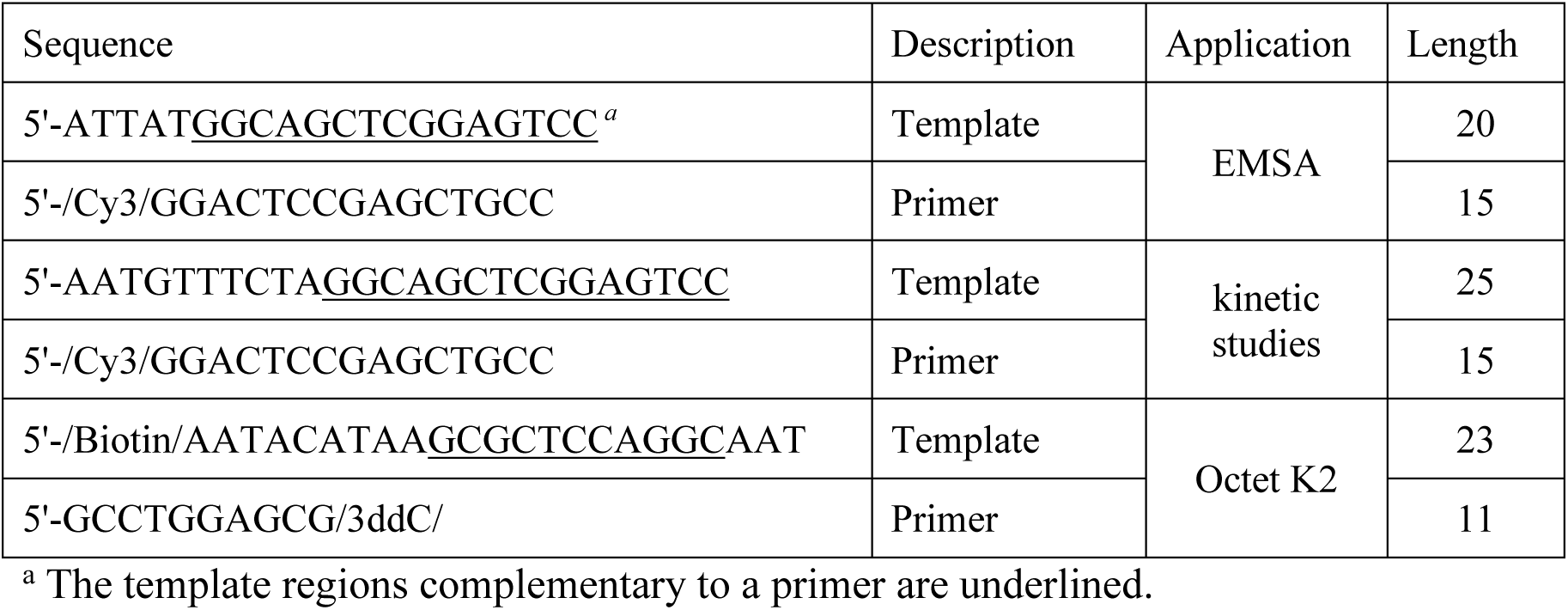
Oligonucleotides used in this study.

**Figure 2.**
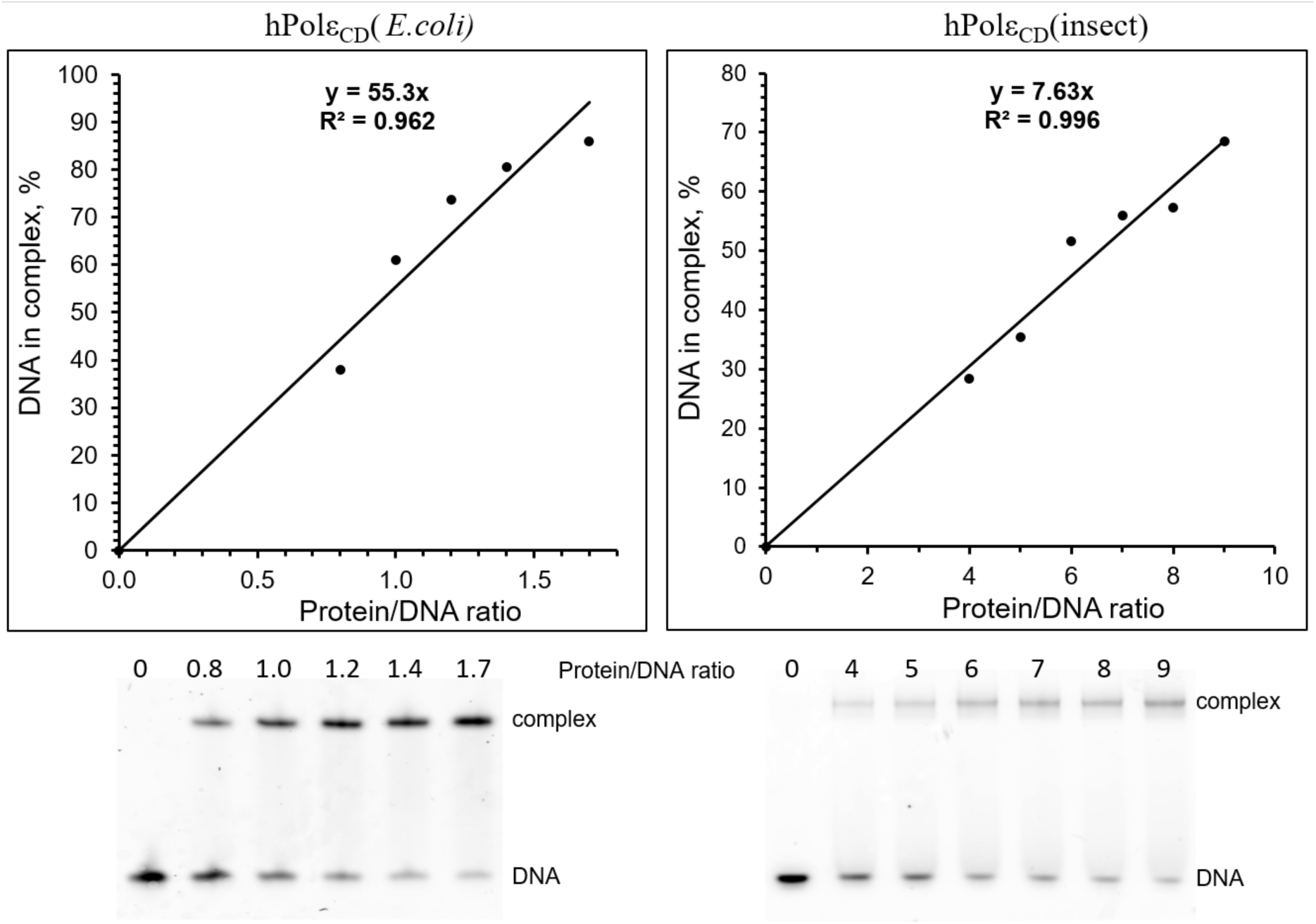
Analysis of the level of active molecules by EMSA. The Cy3-labeled DNA (0.5 μM) was incubated with varying amount of protein. Samples were separated by electrophoresis in 5% acrylamide gel and visualized by Typhoon FLA 9500. The percent of DNA in the complex is plotted against the protein/DNA ratio, and the generated trend line shows the percent of active molecules.

### hPolε_CD_ mutant deficient in the [4Fe-4S] cluster displays no DNA interaction and is inactive

In order to confirm that the [4Fe-4S] cluster is required for interaction with a template:primer, we obtained the hPolε_CD_ mutant (hPolε_CD_^M^) where two of four conserved cysteines (654 and 663) coordinating the cluster were mutated to serines. Interestingly, upon elution from a Heparin HiTrap HP column by a gradient of sodium chloride, the mutant was eluted at a lower salt concentration than the intact hPolε_CD_ (SI Fig. S1). Heparin HiTrap is considered an affinity column for DNA-binding proteins because its sulfate groups mimic phosphates of DNA, so the earlier mutant elution from it indicates compromised interaction with DNA. hPolε_CD_^M^ is contaminated with a proteolysis product of a molecular mass of ∼120 kDa, a likely cause of its lower affinity to the Heparin column. Purification of hPolε_CD_^M^ by a size-exclusion column revealed significant level of aggregates that were eluted in the void volume (SI Figure S2). Of note, a slightly increased tendency to aggregate was mentioned for yeast Polε_CD_ mutant deficient in the [4Fe-4S] cluster (10). The concentrated hPolε^CD^_M_ sample was colorless and iron-free as was previously shown for yeast Polε mutants (10,11).

Analysis of DNA-binding activity by EMSA revealed that interaction with DNA is significantly compromised in the case of a mutant (Figure 3). Even at four-fold molar excess over DNA, hPolε_CD_^M^ had no effect on mobility of single- and double-stranded DNA. In contrast, hPolε_CD_ showed a good shift for both DNA substrates. Next, we analyzed the DNA polymerase and exonuclease activities of hPolε_CD_ and the mutant using a Cy3-labeled 15-mer primer annealed to a 25-mer template. Both activities require the presence of magnesium ions. In addition, a dNTP mix was added to the DNA polymerizing reactions. In contrast to hPolε_CD_, the mutant showed no activity in either assay (Figure 4). Thus, the absence of a [4Fe-4S] cluster severely affects DNA-binding, DNA-polymerase, and exonuclease activities of hPolε on the primed template. Interestingly, the mutant showed good exonuclease activity on the primer alone (Figure 5). These data indicate that single-stranded DNA is able to reach the exonuclease active site of hPolε_CD_^M^.

**Figure 3.**
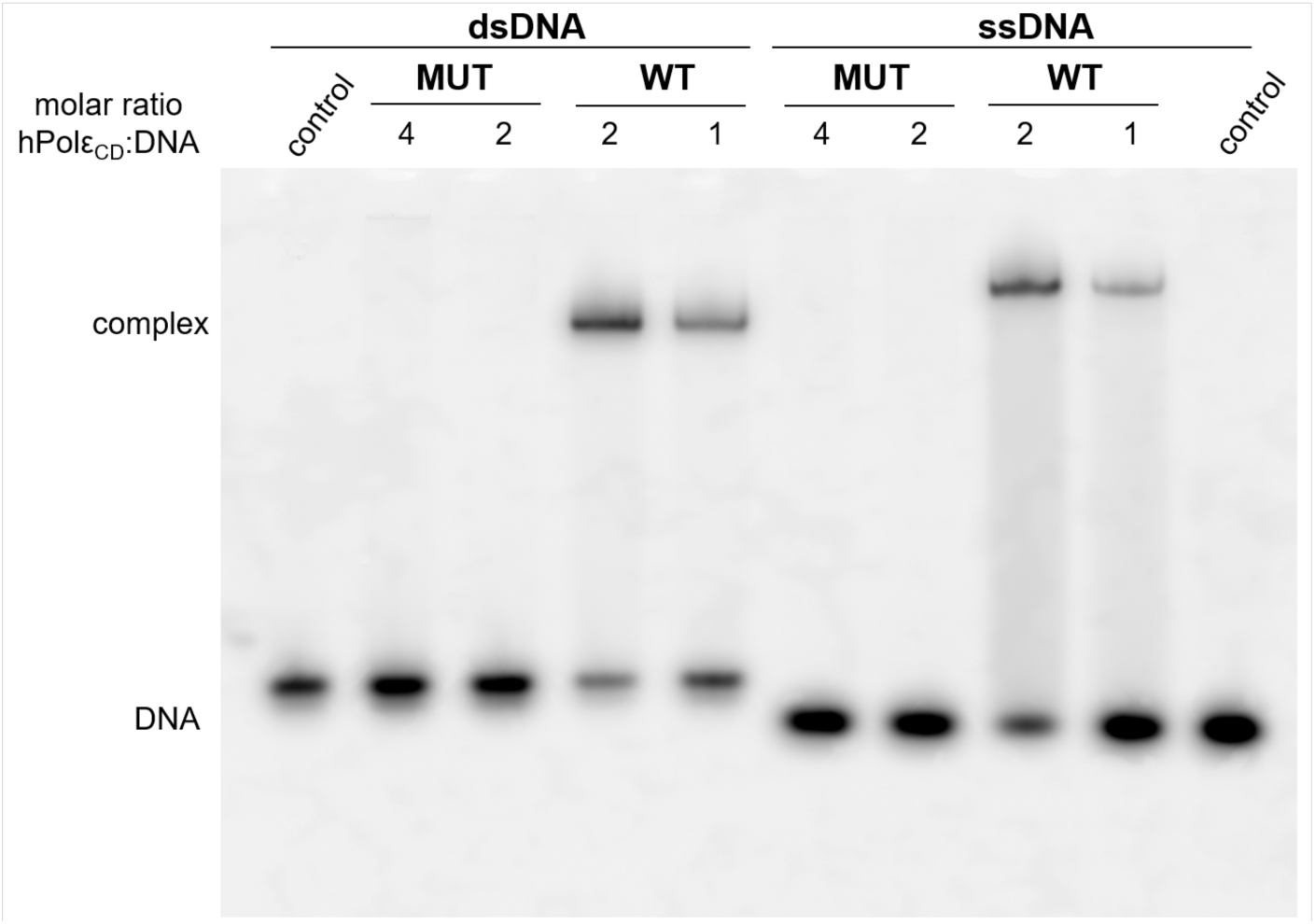
hPolε_CD_ mutant does not bind DNA. A 15-mer Cy3-labeled primer alone (ssDNA) or in the duplex with a 25-mer template (dsDNA) was incubated for 5 min with hPolε_CD_ and its mutant. Samples were separated by electrophoresis in 5% acrylamide gel and visualized by Typhoon FLA 9500.

**Figure 4.**
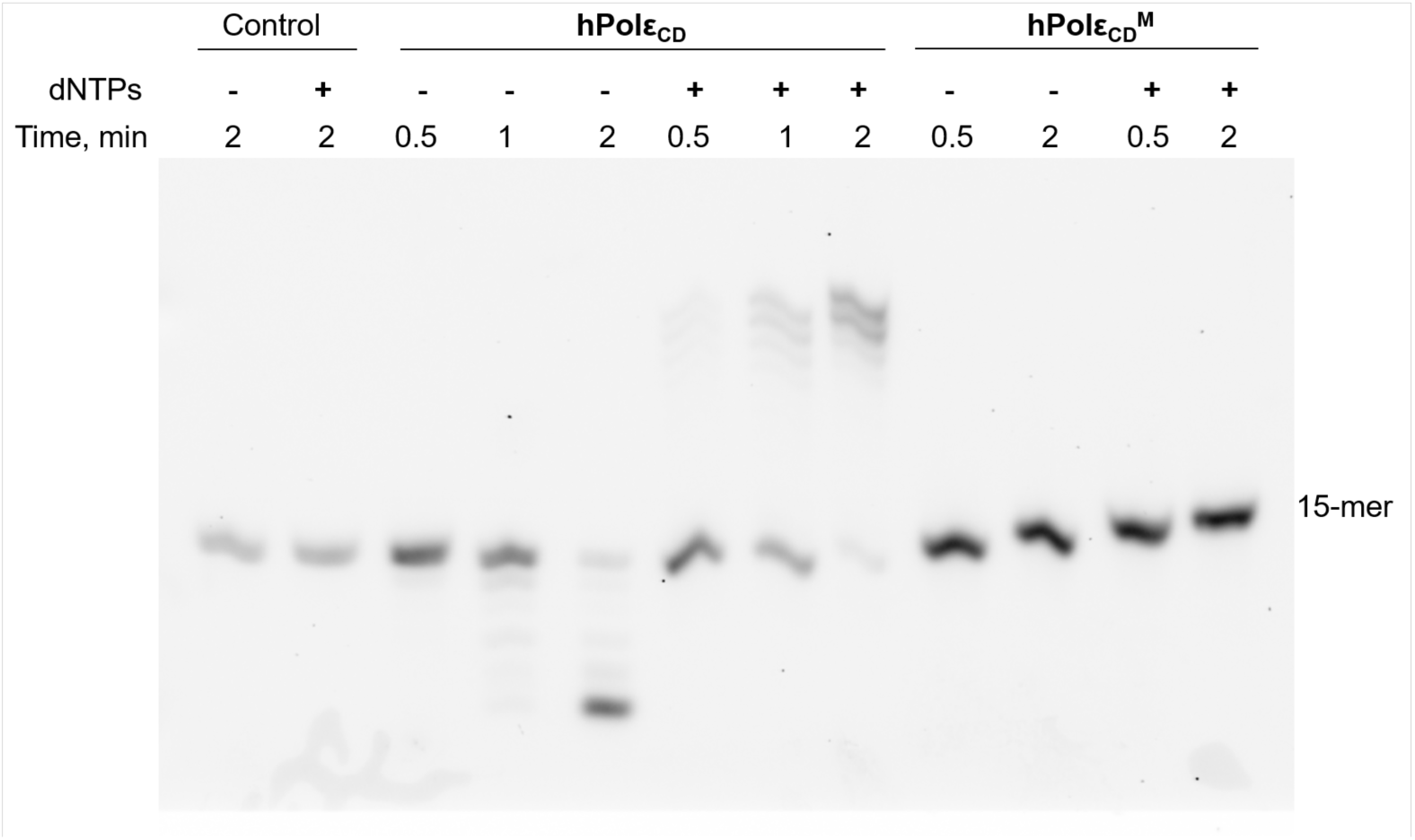
hPolε_CD_^M^ does not exhibit DNA-polymerase and exonuclease activities on the primed template. Primer extension assay was performed at 35ºC in the presence of dNTPs and a DNA substrate with a Cy3-labeled 15-mer primer annealed to a 25-mer template. Exonuclease activity was analyzed in the absence of dNTPs.

**Figure 5.**
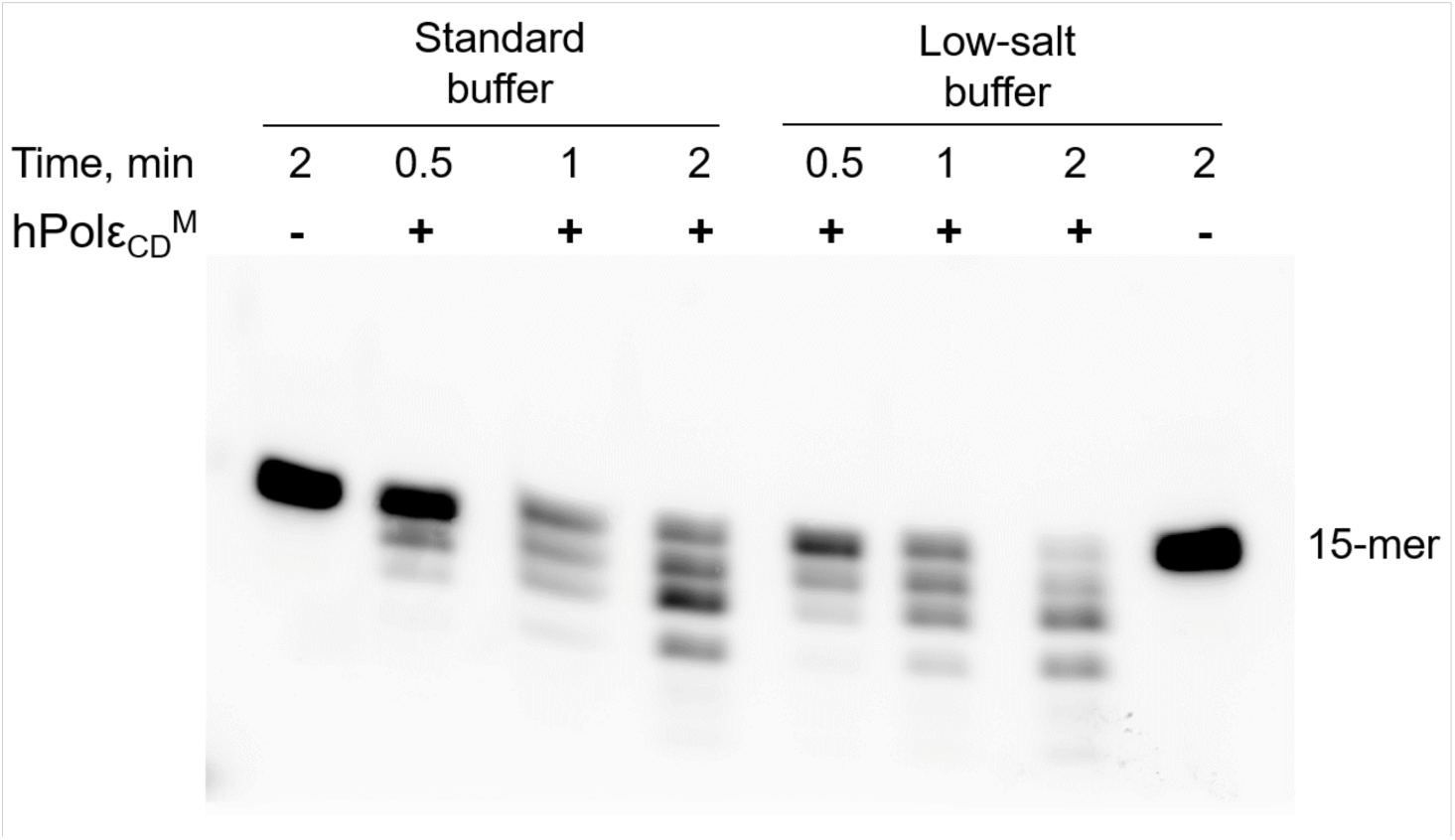
hPolε_CD_^M^ exhibits exonuclease activity on single-stranded DNA. Exonuclease assay was performed at 35ºC using a 15-mer single-stranded DNA labeled with a Cy3 fluorophore at the 5’-end (0.2 μM); hPolε_CD_^M^ concentration in reaction was 0.1 μM. The low salt buffer contains 25 mM Tris-HEPES, pH 7.8, 40 mM NaAc, 8 mM Mg(Ac)_2_, 2 mM TCEP, and 0.2 mg/mL BSA.

### Binding and kinetic studies of Polε_CD_

The analysis of Polε_CD_ interaction with DNA was conducted using an Octet K2, which allows for extraction of the rate constants of complex formation (*k*_on_), dissociation (*k*_off_), and the dissociation constant (*K*_*D*_), which is inversely proportional to affinity. A 23-mer DNA template with biotin at the 5’-end (Table 2) was primed by an 11-mer DNA primer and loaded on a streptavidin-coated sensor. In the presence of 0.1 M NaCl, hPolε_CD_ samples obtained from *E. coli* and insect cells have shown similar *K*_*D*_ values close to 200 nM (Table 3). A previous DNA-binding study of hPolε_CD_ conducted in the absence of salt reported a *K*_*D*_ value of 79 nM (16). Notably, an increase of NaCl concentration from 0.1 M to 0.15 M reduced affinity ∼20-fold, which is mainly due to the 13-fold reduction in *k*_off_ value (Table 3). We observed a similar effect of salt concentration on affinity to DNA for Polα_CD_ (17) and for a different hPolε_CD_ construct, deficient in exonuclease activity (18). Thus, hPolε_CD_ binds a DNA duplex with relatively low affinity at near-physiological salt concentration (Table 3, *K*_*D*_ = 3.3 μM). The obtained *k*_off_ value of 0.54 s^-1^ indicates that the half-life of the hPolε_CD_/DNA complex is ∼1.3 s on average. It was not possible to conduct binding studies for hPolε_CD_(insect) at 0.15 M NaCl with acceptable accuracy due to the low level of active molecules, which demands very high protein concentration in reaction.

**Table 3.** Interaction of hPolε_CD_ with DNA.

| Expression system | [NaCl]<br>mM | $k_{on}$<br>mM<sup>-1</sup> sec<sup>-1</sup> | $k_{off}$ x10<sup>-3</sup><br>sec<sup>-1</sup> | $K_D$ <sup>a</sup><br>nM |
| --- | --- | --- | --- | --- |
| insect | 100 | 162 ± 2.5 | 34.2 ± 6.3 | 213 ± 43 |
| *E. coli* | 100 | 252 ± 26 | 41.9 ± 6.7 | 167 ± 24 |
| *E. coli* | 150 | 168 ± 29 | 542 ± 15 | 3258 ± 386 |

<sup>a</sup> $K_d$ values are obtained by dividing $k_{off}$ by $k_{on}$ .
Data are presented as mean ± SD.

The pre-steady-state kinetic approach was employed in order to assess hPolε_CD_ activity in DNA primer extension and its affinity to incoming dNTP at conditions used for binding studies. Kinetic studies were conducted in the presence of 0.1 M NaCl to allow most of the DNA to complex with hPolε_CD_ before the reaction. Single-nucleotide incorporation experiments were conducted under single-turnover conditions. This assay provides the maximal polymerization rate (*k*_pol_) and the apparent dissociation constant (*K*_*D*_) for the incoming nucleotide. hPolε_CD_ (0.8 μM) was incubated with a Cy3-labeled DNA (0.4 μM) and quickly mixed with varying dTTP concentrations under rapid chemical quench conditions. For each dTTP concentration, the fraction of extended primer was plotted against time (Figure 6*A*) and the data were fit to a single-exponential equation (Eq. 1).

**Figure 6.**
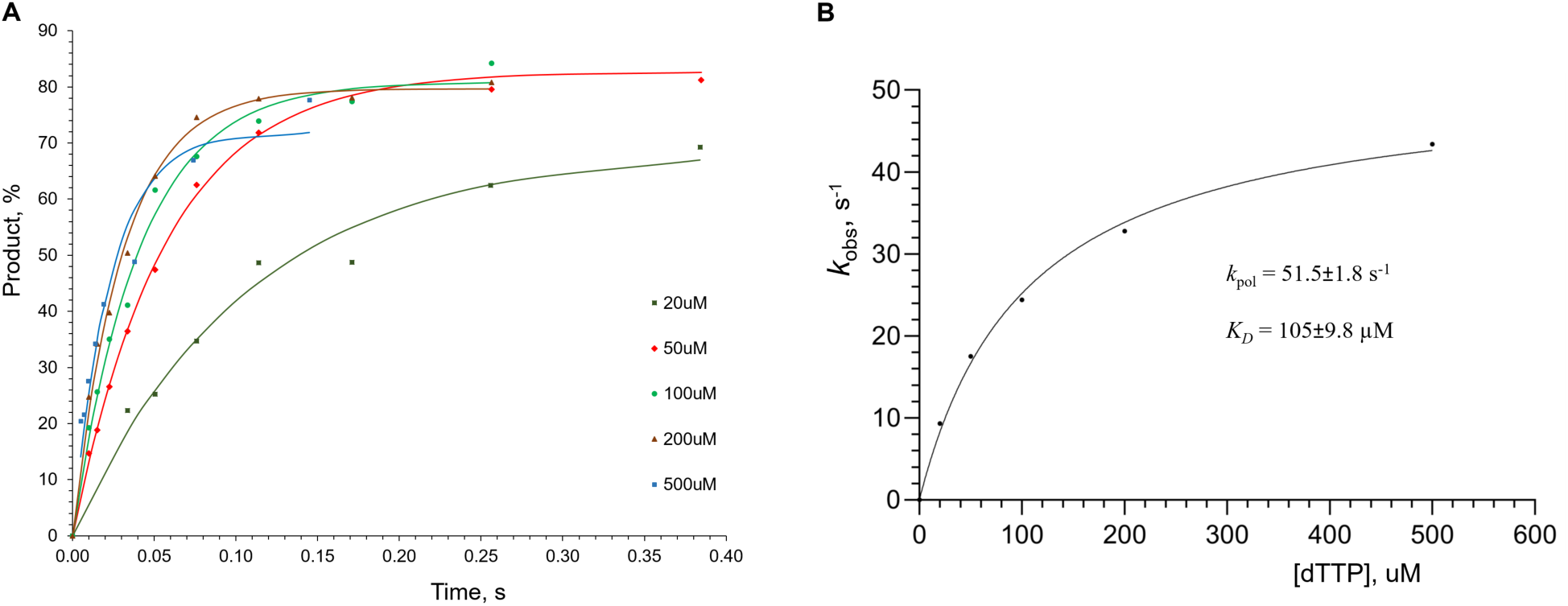
Single turnover kinetics of primer extension by hPolε_CD_. *A*, Percent of extended primer was plotted against time and the data were fit to a single-exponential equation (Eq. 1). *B*, Primer extension rates are plotted against dTTP concentration and the data were fit to a hyperbolic equation (Eq. 2) to obtain *K*_*D*_ and *k*_pol_ values. Reactions, containing 0.8 μM hPolε_CD_, 0.2 μM DNA, and dTTP at varied concentrations, were incubated at 35ºC at indicated time points.

The obtained rate constant values were plotted against dTTP concentration (Figure 6*B*) and fit to (Eq. 2), resulting in a *k*_pol_ of 51.5±1.8 s^-1^ and an apparent *K*_*D*_ of 105±9.8 μM. A *K*_*D*_ value of 31 μM was reported previously for the hPolε_CD_(exo-)/dTTP complex by another group (16). The 3.4-fold difference with our data might be attributable to the absence of salt in reaction in that study. As shown above (Table 3), salt significantly affects the hPolε_CD_/DNA complex. The same effect is expected for the hPolε_CD_/dNTP complex because their interaction interface is mainly electrostatic. An apparent *K*_*D*_ of 105 μM for dTTP indicates that hPolε might not be saturated with dNTPs *in vivo* to achieve the maximal polymerization rate, given the fact that the average concentration of each dNTP in human mitotic cells is approximately 50 μM (19) or even lower (20). On the other hand, it is possible that the local concentration of dNTPs near the replication fork is significantly elevated. hPolε_CD_(insect) was not suitable for this assay because of its low level of active molecules. In another study, we have shown that the exonuclease deficient variant of hPolε_CD_ extends the primer at a rate of 62.4 s^-1^ in the presence of 1 mM dTTP (18). Interestingly, a *k*_pol_ value of 248 s^-1^ was obtained for hPolε_CD_ at 20 ºC (16), which might be due to the absence of salt in reaction (21). Noteworthy, the catalytic domain of human Polα showed the maximal rate of DNA polymerization of 33.8 s^-1^ in the presence of 0.1 M NaCl (22).

## DISCUSSION

The recently discovered [4Fe-4S] cluster, located at the base of the P-domain (Figure 7) and coordinated by four conserved cysteines, is unique to Polε (10,11). Mutation of these cysteines in yeast Polε led to a loss of iron and DNA polymerase activity but not exonuclease activity, as was shown for holoenzyme as well as a separate catalytic domain (10,11). The role of the [4Fe-4S] cluster in the DNA-binding properties of Polε has not been studied so far. Our data indicate that the iron-sulfur cluster is critical for interaction of human Polε with DNA. Probably, abrogated DNA binding is responsible for the loss of both activities in hPolε_CD_. Substrate binding by any enzyme is an initial step that precedes catalysis. It is difficult to imagine how the absence of the distantly located iron-sulfur cluster can affect both catalytic centers and especially exonuclease. Structural data indicate that the [4Fe-4S] cluster may stabilize the P-domain (Figure 7) (11). It is possible that in the absence of the cluster, the P-domain slightly changes its position and blocks entrance into the DNA-binding pocket for a template:primer.

**Figure 7.**
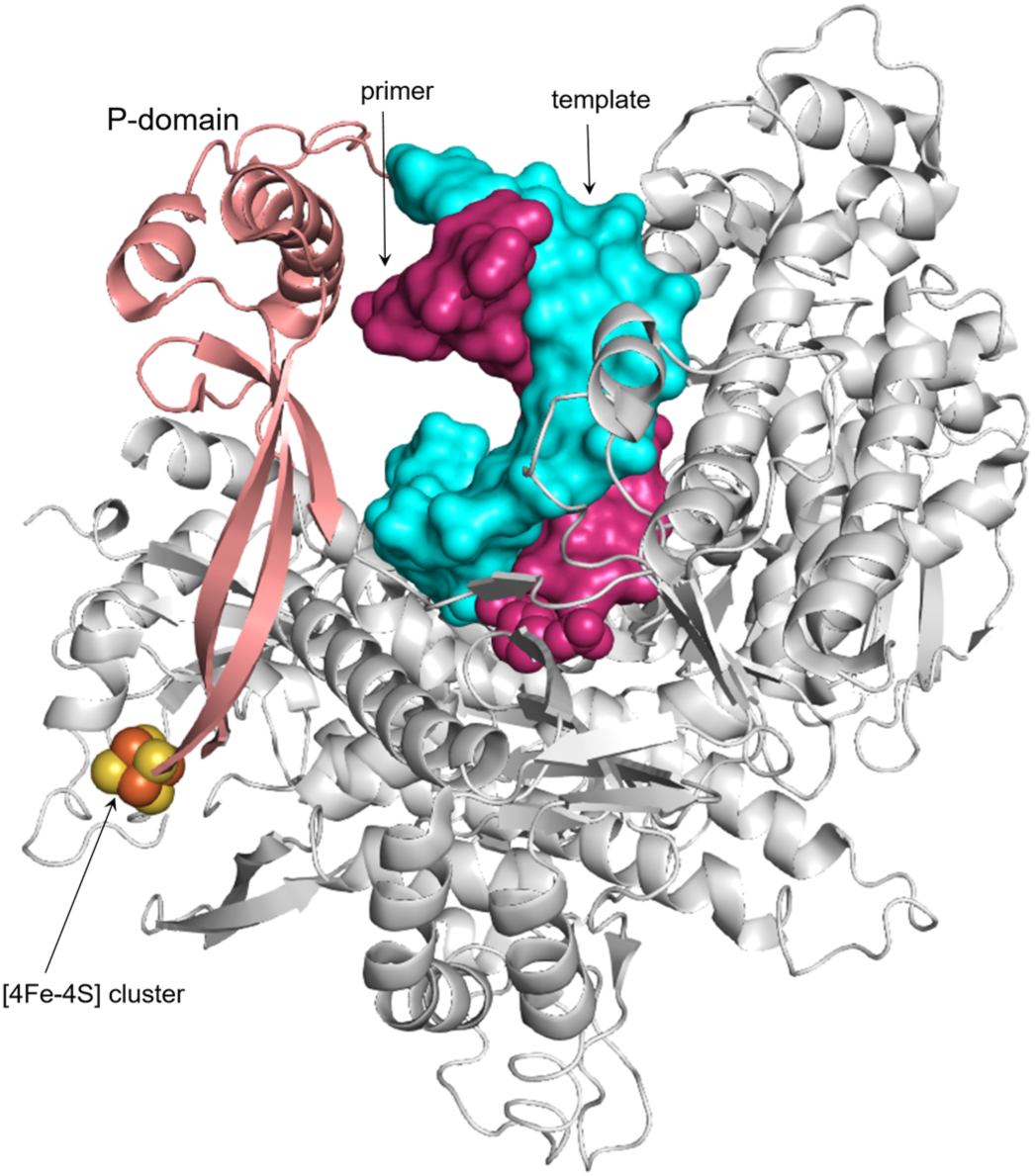
Position of the P-domain and the iron-sulfur cluster in yeast Polε. The P- and thumb domains and the rest of a protein are colored salmon, green, and gray, respectively. Template and primer are presented as surface and colored as cyan and pink, respectively. The 3’-terminal nucleotide of a primer is presented as spheres. The iron and sulfur in the [4Fe-4S] cluster are presented as spheres and colored brown and yellow, respectively. The crystal structure of a ternary complex of yeast Polε_CD_ with DNA and dATP (pdb code 6QIB (22)) was used for this presentation.

It is quite possible that inactivation of DNA polymerase activity of yeast Polε by mutation of cluster-coordinating cysteines is also due to abrogated DNA binding and not catalysis. Intriguingly, the exonuclease activity of yeast Polε on the primed template was not affected upon disruption of the [4Fe-4S] cluster (10,11). Moreover, DNA polymerase activity slightly recovered when dNTPs concentration was increased ten-fold, pointing to compromised dNTP binding (11). We cannot exclude possibility that elimination of the cluster has a slightly different effect on the structure and activity of yeast and human Polε. For example, in cluster-deficient mutants of yeast Polε, the template:primer-binding cleft might be blocked at smaller degree than in hPolε_CD_^M^, Of note, in reactions with yeast Polε mutants, DNA substrates were added at tenfold lower concentration and had higher AT-content at the primer 3’-end in comparison to our studies, so the DNA duplex might be partially melted at 30 ºC providing single-stranded DNA for exonuclease.

The biological role of the [4Fe-4S] cluster in regulating DNA-binding properties and activity of Polε requires further investigation. As previously proposed (10), due to its peripheral position and potential sensitivity to oxidation, the cluster might be a target for oxygen reactive species during oxidative stress. Recently it was demonstrated that the [4Fe-4S] cluster of yeast Polε is redox active and its reversible oxidation affects DNA polymerase activity (15). We propose that the main role of the P-domain in hPolε is controlling access to the DNA-binding pocket, which might depend on intracellular signals. Even a subtle change in the P-domain position would be enough to prevent template:primer binding (Figure 7).

This work revealed that insect cells do not efficiently incorporate the [4Fe-4S] cluster into heterologous proteins during overexpression. This may be due to low capacity and/or high specificity of the enzymatic machinery, which is responsible for cluster incorporation. In contrast, the corresponding machinery of bacteria is more robust and non-specific. Previously we have shown that the [4Fe-4S] cluster was erroneously incorporated at a significant level into a zinc-binding domain located at the C-terminus of the catalytic subunit of hPolε (13,14).

Fortunately, the cluster-containing molecules were unable to make a stable complex with an accessory subunit of hPolε and were separated during purification. Thus, only the Polε catalytic domain contains the iron-sulfur cluster (10,11). This study underscores the importance of measuring the iron level in Polε samples, especially when the enzyme is overexpressed in insect cells. Notably, all previous functional studies of hPolε_CD_ were conducted with a recombinant protein expressed in *E. coli*, where the iron content was not analyzed (16,23-25).

## MATERIALS AND METHODS

### Cloning, expression, and purification

hPolε_CD_ was cloned into pASHSUL-1 plasmid (26) to produce the corresponding proteins tagged with N-terminal His-Sumo. The mutant with cysteines 654 and 663 changed to serines was obtained by side-directed mutagenesis. hPolε_CD_(insect) with a cleavable N-terminal His-Tev tag was cloned into pFastBac-1 plasmid (Invitrogen). hPolε_CD_ and hPolε_CD_(exo-) were expressed in *E. coli* strain Rosetta-2 (DE3) at 18 ºC for 16 h following induction with 0.2 μg/ml anhydrotetracycline. Afterward, cells were harvested by centrifugation at 4,000 g for 15 min, washed with PBS, aliquoted, and kept at -80 ºC. A high-titer virus stock for hPolε_CD_(insect) was obtained by using the Bac-to-Bac Baculovirus Expression System from Invitrogen. 1.8×10^9^ Sf21 cells in 1 L shaking culture were infected with the recombinant virus at a multiplicity of infection of 2 and cultivated at 27 ºC for 56 hr. Cells were harvested by centrifugation at 200 g for 5 min and frozen.

All samples were purified according to the same protocol, including chromatography on a Ni-IDA column (Bio-Rad), His-tag digestion during overnight dialysis, and chromatography on a Heparin HP HiTrap column (Cytiva) and on a size-exclusion column Superose 12 10/300 GL (Cytiva) in buffer containing 25 mM Tris-HEPES (pH 7.8), 0.15 M NaCl, 1% glycerol, and 2 mM tris(2-carboxyethyl)phosphine (TCEP). Finally, samples were concentrated to 30–60 μM and flash-frozen in aliquots. The purity of obtained samples was analyzed by 8% SDS-PAGE (Figure 1). Protein concentrations were estimated by measuring the absorbance at 280 nm and using extinction coefficients of 157 mM^-1^ cm^-1^; the extinction coefficients were calculated with ProtParam (27). The iron content in purified protein samples was determined with use of chromogen ferrozine as described in (22).

### Electrophoretic mobility shift assay

Reactions containing 0.5 μM DNA (Table 2) and varying amount of protein were incubated in 10 μl for 5 minutes at room temperature in buffer containing 20 mM Tris-Hepes (pH 7.8), 100 mM NaCl, 2% glycerol, 2 mM TCEP, and 0.2 mg/mL BSA; 5μl was then loaded on 5% native PAGE. Samples labeled with Cy3-dye were visualized using a Typhoon 9410 imager (Cytiva) and quantified using ImageJ software (version 1.45s, National Institutes of Health).

### Binding studies

Analysis of binding kinetics was done at 23 ºC on an Octet K2 (Sartorius AG). This device uses Bio-Layer Interferometry technology to monitor molecular interactions in real time. A template with a biotin-TEG at the 5’-end was annealed to the primers (Table 2) and immobilized on a streptavidin-coated biosensor (SAX, Sartorius AG). Primers were added at two-fold molar excess over the template. SAX sensors were loaded with oligonucleotide-biotin at 50 nM concentration for 7 min at 500 rpm. Then sensors were blocked by incubating for 2 min in 10 μg/ml biocytin. In the first row of a 96-well microplate (Greiner Bio-One), the first six wells contained the buffer, consisting of 30 mM Tris-Hepes, pH 7.8, 100 mM or 150 mM NaCl, 2 mM TCEP, and 0.002% Tween 20. The next six wells contained the two-fold dilutions of hPolε_CD_ in the same buffer. All wells in the adjacent row contained only the buffer for reference. Data Analysis HT software (ver. 11.1, Sartorius AG) was used for calculation of binding constants (*k*_on_, *k*_off_, and *K*_*D*_) by using the global fitting. The average value and standard deviation were calculated from three independent experiments.

### Pre-steady-state kinetic studies

Kinetic studies were performed on a QFM-4000 rapid chemical quench apparatus (BioLogic, France) at 35 °C. Reactions contained 0.4 μM hPolε_CD_ (active molecules), 0.2 μM DNA, varying concentrations of dTTP, 25 mM Tris-HEPES, pH 7.8, 0.1 M NaCl, 8 mM MgCl_2_, 2 mM TCEP, and 0.2 mg/mL BSA. hPolε_CD_ was incubated with a Cy3-labeled 15-mer primer annealed to a 25-mer DNA template (Table 2), to allow formation of the binary complex, and rapidly mixed with dTTP and MgCl_2_ followed by quenching with 0.3 M EDTA. Products were collected in a tube containing 15ul 100% formamide and separated by 20% Urea-PAGE. The Cy3-labeled products were visualized by a Typhoon FLA 9500 (Cytiva) and quantified by ImageJ, version 1.5.3 (NIH). The extended primer fraction was calculated by dividing the amount of extended primer by the amount of primer added in reaction. For each dTTP concentration, the percent of extended primer was plotted against time and the data were fit to a single exponential equation:

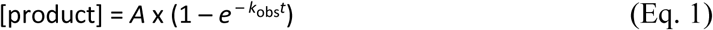

where *A* is the amplitude, *k*_*obs*_ is the observed rate for dNTP incorporation, and *t* is the time. The

*k*_*obs*_ was plotted against dTTP concentration and the data were fit to the hyperbolic equation:

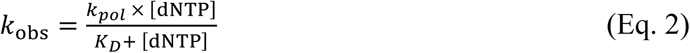

using GraphPad Prizm software to obtain *k*_*pol*_, the maximum rate of nucleotide incorporation, and *K*_*D*_, the apparent dissociation constant for the incoming nucleotide.

### DNA polymerase and exonuclease assay

Exonuclease reactions were conducted in 10 μl at 35 °C and contained 0.05 μM hPolε_CD_ or its mutant (total protein concentration), 0.2 μM DNA (Cy3-labeled 15-mer primer annealed to a 25-mer DNA template), 25 mM Tris-HEPES, pH 7.8, 0.1 M NaCl, 5 mM MgCl_2_, 2 mM TCEP, and 0.2 mg/mL BSA. DNA polymerase reactions were performed in a similar way except using 10 μM protein and adding 50 μM dNTPs. The reactions were stopped by addition of 20 μl stop solution (96% formamide, 50 mM EDTA, 0.1% bromophenol blue) and heated to 95 ºC for 1 min. The products were separated on a 20% denaturing polyacrylamide gel and visualized by a Typhoon FLA 9500 (Cytiva).

This article contains supporting information

## Acknowledgements

We thank K. Jordan for editing this manuscript.

## Author contributions

A.E.L., A.G.B., and L.M.M. purified hPolε_CD_ samples. A.E.L. performed functional studies. L.M.M., N.D.B., and E.I.S. prepared the plasmids encoding different hPolε constructs and analyzed their expression. A.G.B. and T.H.T. initiated and supervised the project. A.G.B. and A.E.L. wrote the manuscript, with contributions and critical comments from the other authors.

## Funding and additional information

This work was supported by the National Institute of General Medical Sciences (NIGMS) grant R35 GM127085 to THT. The University of Nebraska Medical Center (UNMC) Genomics Core receives partial support from the National Institute for General Medical Science (NIGMS) INBRE - P20GM103427 grant as well as the Fred & Pamela Buffett Cancer Center Support Grant - P30 CA036727. The content is solely the responsibility of the authors and does not necessarily represent the official views of the National Institutes of Health.

## Abbreviations

hPolε: human DNA polymerase ε
hPolε_CD_: catalytic domain of human DNA polymerase ε
P-domain: processivity domain
EMSA: electrophoretic mobility gel shift assay
TCEP: tris(2-carboxyethyl)phosphine.

