## Supplemental Figures 1 and 2 for "The iron-sulfur cluster is critical for DNA binding by human DNA polymerase ε"

**for the article**

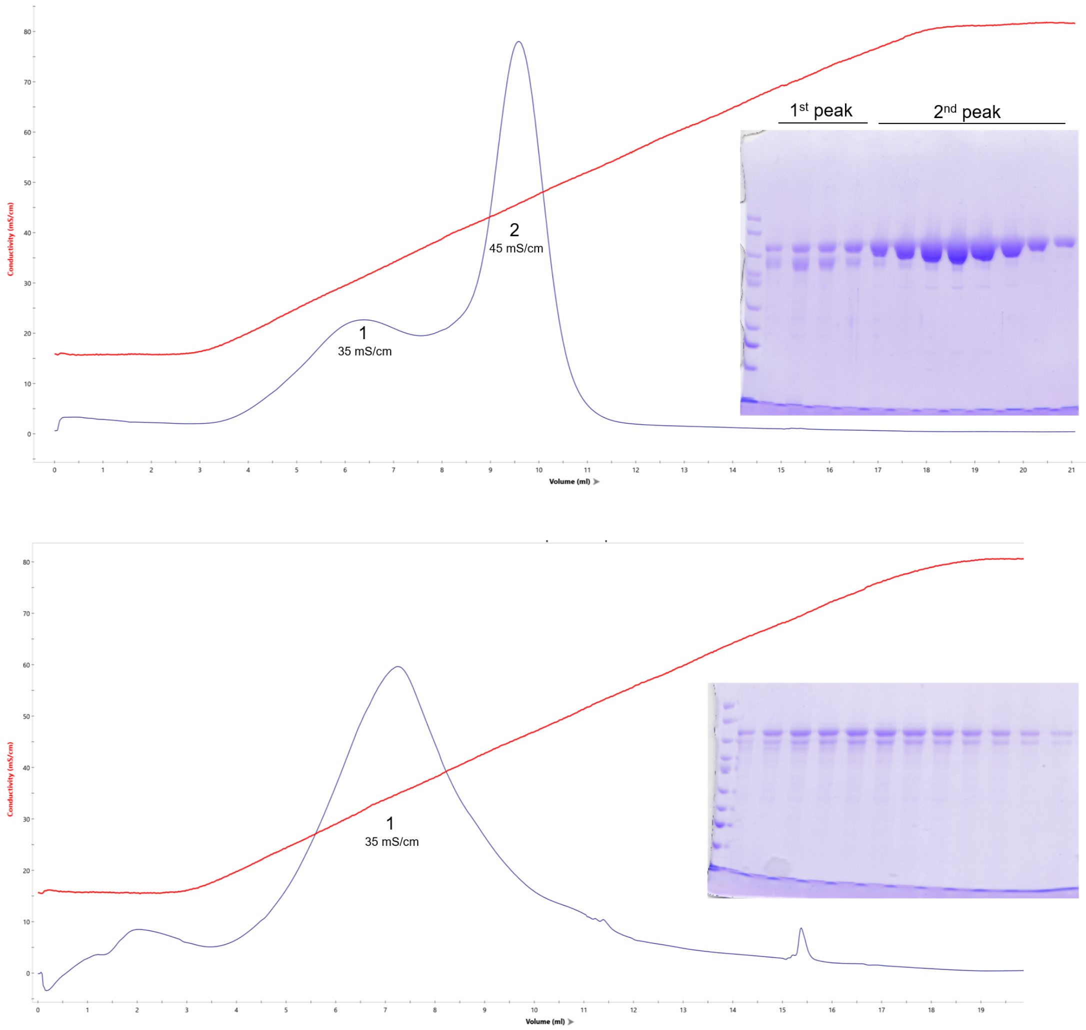


**Figure S1. Heparin elution profiles of hPol$\varepsilon_{CD}$ (A) and hPol$\varepsilon_{CD}^{M}$ (B).** Heparin HiTrap HP column (Cytiva, 1 ml) was eluted by 15 ml gradient of 0.15 - 0.8 M NaCl. Peak fractions were analyzed by 8% SDS-PAGE and stained by Coomassie Brilliant Blue R-250. The mutant elutes as a wide peak and at lower salt concentration than wild-type hPol$\varepsilon_{CD}$, resulting in contamination with a proteolyzed form.

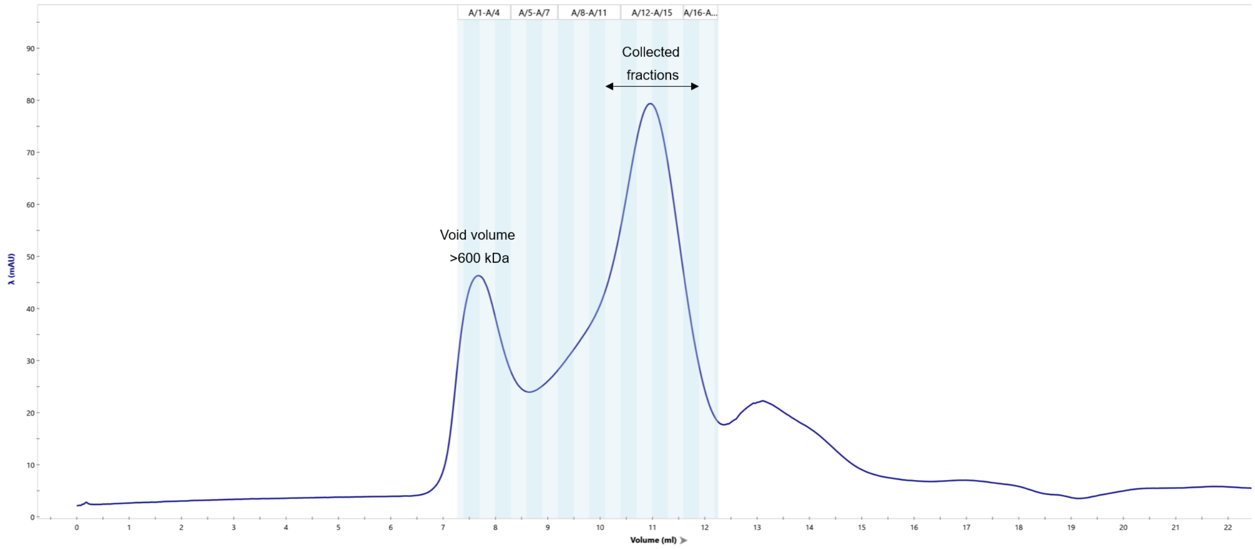


**Figure S2. Elution profile of hPolε$_{CD}$$^{M}$ on Superose 12 size exclusion column.** Superose 12 10/300 GL (Cytiva) has a bed volume of 24 ml and an exclusion limit of 300 kDa for globular proteins. The chromatography was conducted at 4ºC in the buffer containing 25 mM Tris-HEPES (pH 7.8), 0.15 M NaCl, 1% glycerol, and 2 mM TCEP.
